# NovoBoard: a comprehensive framework for evaluating the false discovery rate and accuracy of de novo peptide sequencing

**DOI:** 10.1101/2024.04.16.589668

**Authors:** Ngoc Hieu Tran, Rui Qiao, Zeping Mao, Shengying Pan, Qing Zhang, Wenting Li, Lei Xin, Ming Li, Baozhen Shan

## Abstract

De novo peptide sequencing is a fundamental research area in mass spectrometry (MS) based proteomics. However, those methods have often been evaluated using a couple of simple metrics that do not fully reflect their overall performance. Moreover, there has not been an established method to estimate the false discovery rate (FDR) and the significance of de novo peptide-spectrum matches (PSMs). Here we propose NovoBoard, a comprehensive framework to evaluate the performance of de novo peptide sequencing methods. The framework consists of diverse benchmark datasets (including tryptic, nontryptic, immunopeptidomics, and different species), and a standard set of accuracy metrics to evaluate the fragment ions, amino acids, and peptides of the de novo results. More importantly, a new approach is designed to evaluate de novo peptide sequencing methods on target-decoy spectra and to estimate their FDRs. Our results thoroughly reveal the strengths and weaknesses of different de novo peptide sequencing methods, and how their performances depend on specific applications and the types of data. Our FDR estimation also shows that some tools may perform better than the others in distinguishing between de novo PSMs and random matches, and can be used to assess the significance of de novo PSMs.

## Introduction

De novo peptide sequencing is essential for the identification of novel peptides or proteins that do not exist in any databases, such as mutated neoantigens or antibody variable regions^1^. This has been an intensive research area in mass spectrometry (MS) based proteomics, with dozens of algorithms proposed over the last three decades^2–5^. However, a fundamental problem, but often overlooked or simplified, is how to evaluate and benchmark the performance of de novo peptide sequencing. For instance, despite having a long history as the database search approach, de novo peptide sequencing still does not have a standard method to estimate the false discovery rate (FDR), as opposed to its counterpart. In the database search approach, a database peptide is a real peptide and we only need to assess the significance of its peptide-spectrum match (PSM) compared to a random match. However, a de novo peptide may still contain sequence errors even when its PSM has a very high score. Thus, FDR estimation is challenging for de novo peptide sequencing, but at least one could attempt to obtain a lower bound approximation and to evaluate the significance of de novo PSMs.

The performance evaluation has recently become more critical due to the rise of new de novo sequencing algorithms that were based on deep learning and large-language models (LLMs). Given the large scale of such models and the massive amount of their training data, inherent bias is inevitable, and the question is how to ensure that a model can truly discover new peptides rather than just memorizing millions of peptides and spectra from its training. Thus, a comprehensive framework with carefully designed metrics to evaluate the performance of de novo peptide sequencing is highly desired, now more than ever.

While the goal is to identify unknown peptides, de novo peptide sequencing methods have traditionally been evaluated on well-studied samples with known ground-truth peptides. Evaluation datasets often consist of peptide-spectrum matches (PSMs) obtained from the database search results of common organisms such as human, mouse, yeast, etc. Database peptides identified at strict FDR levels of 1%, or 0.1%, are considered as ground-truth to evaluate the de novo peptides identified on the same spectra. However, such an evaluation approach suffers from several major limitations. Firstly, the evaluation spectra identified at those strict FDR levels are more likely to have very good fragmentations and high signal-to-noise ratios. They often account for less than 20-30% and are not a good representation of the whole dataset, especially for spectra of medium to low quality where unknown peptides may come from. Secondly, the evaluation peptides are also highly biased because they come from well-known databases that have been analyzed numerous times. Thus, the peptides and spectra obtained from the database search may be very different from the unknown peptides and spectra that a de novo sequencing algorithm is supposed to find. As a result, the current evaluation approach may not accurately capture the real performance of de novo peptide sequencing.

The third limitation arises from recent deep learning models developed over the last few years. Those models are purposely designed to learn much more patterns of fragment ions and amino acid sequences to predict more accurate peptides. However, it is also their impressive learning capability that may lead to overfitting and memorizing, rather than generalizing and discovering new sequences. It is not uncommon to observe that some deep learning algorithms, particularly the ones utilizing peptide language models, can predict peptide sequences with high-confidence scores while their spectra have very few supporting fragment ions. Some efforts have been put into controlling the overfitting issue, such as splitting the data into training and testing sets with non-overlapping peptide sequences, or using two different species for training and testing.

However, due the nature of high similarity of genomic and protein sequences, two different proteins or peptides may still share sub-sequences. Thus, information from the training may be leaked to the evaluation. It is not trivial how to address this problem, especially when the models and the data are getting larger and larger.

## Results

To overcome the aforementioned limitations, here we propose NovoBoard, a new framework that includes benchmark datasets and metrics to evaluate the performance of de novo peptide sequencing methods.

### Benchmark Datasets

First, we carefully selected five different types of benchmark datasets that represent a wide range of applications of de novo peptide sequencing. The first one is a standard human tryptic dataset from the 2017 Association of Biomolecular Resource Facilities (ABRF) Proteomics Research Group (PRG) study. This is a typical MS dataset where most de novo sequencing methods should achieve very good performance, if not the best performance among the five datasets. The second one is a nontryptic dataset from Wang et al.^6^ (PXD010154) that includes six enzymes: Arg-C, Asp-N, Chymotrypsin, Glu-C, Lys-C, and Lys-N. Since many de novo sequencing models have been trained mainly on tryptic data, it is important to assess how they would perform on nontryptic data. We also selected a dataset from *Arabidopsis thaliana* (*A. thaliana*, PXD013658, Muntel et al.^7^), a species that is less related to human and other common model organisms. To the best of our knowledge, *A. thaliana* data has not been used to train any de novo sequencing models. Furthermore, we performed evaluation on an HLA class 1 dataset (PXD021013) from Wilhelm et al.^8^). MS-based immunopeptidomics is a major application of de novo peptide sequencing since this technique is essential to identify mutated neoantigens that do not exist in any databases. In addition, due to their unspecific digestion and other unique characteristics such as length and binding motifs, HLA peptides are very different from those peptides derived from trypsin and other enzymes’ digestion. Finally, to evaluate the de novo sequencing methods on unseen mutated peptides, we simulated a dataset of randomly mutated peptides and their *in silico* spectra. First, ten thousand peptides were randomly selected from the MassIVE-KB database^9^, then 1-10 amino acid mutations were introduced to each peptide and finally their spectra were predicted using Prosit^8^.

The raw data of all five datasets were converted to MGF files using the Proteome Discoverer software. We also performed database search of those datasets using Proteome Discoverer with the following parameters: precursor mass tolerance 10 ppm, fragment ion mass tolerance 0.02 Da, fixed Cysteine(Carbamidomethylation), variable Methionine(Oxidation), human and *Arabidopsis thaliana* protein databases (version SwissProt 20240224), and peptide FDR 1%. The MGF files and search results were made publicly available.

### De novo sequencing softwares

In this study we considered four de novo sequencing tools: PEAKS^10^, PointNovo^11^, Casanovo^12,13^, and GraphNovo^14^. However, it should be emphasized that the evaluation framework proposed here can be easily applied to any other tools. In fact, we expect that when more tools are included in the evaluation, new benchmark datasets and metrics will emerge, as different tools may have different strengths and weaknesses.

PEAKS is one of the most common tools for de novo peptide sequencing and has been actively developed and maintained over the last 20 years. In addition, PEAKS is based on a traditional dynamic programming algorithm as opposed to recent deep learning-based methods, thus it could serve as a robust baseline for our evaluation study.

PointNovo and its predecessor DeepNovo^5,15^ were among the first deep learning models developed for de novo peptide sequencing. They have proved that deep learning models are capable of learning considerably more features from the MS data and offer substantial improvements over traditional methods, both in terms of accuracy and methodology. Deep learning models also have the advantages over traditional methods since the data is growing exponentially.

Casanovo model is based on the Transformer^16^ neural network, the popular architecture that powers most of today’s LLMs. The impressive learning capability of Transformer has been demonstrated on large-scale models and data, such as GPT-4^17^, Llama^18^, etc, especially on text data. Thus, one could expect that such a model may also show great performance on protein sequence data, which can be viewed as a special kind of language.

GraphNovo is a recent de novo peptide sequencing model that is based on a graph neural network. It is designed to combine the advantages of both deep learning and graph theory, a popular approach to model the relationships between fragment ions in an MS/MS spectrum^4^. By carefully modeling the features of the fragment ions (nodes) and their relationships (edges) in a spectrum graph, GraphNovo can address the missing-fragmentation problem in de novo peptide sequencing and subsequently improve the sequencing accuracy over previous methods.

### Evaluation metrics for de novo peptide sequencing

To evaluate the results of de novo peptide sequencing, we considered the following four metrics: peptide accuracy, amino acid accuracy, fragment ion accuracy, and an FDR evaluation. The details and motivations of those metrics are explained below.

#### Accuracy metrics for de novo peptide sequencing

Peptide and amino acid accuracies are the two standard evaluation metrics that have been used in most previous studies. In particular, given a tandem mass spectrum and its ground-truth peptide, a de novo peptide predicted from that spectrum is compared to the ground-truth. In addition to exact peptide matching, one can also count how many amino acids in the de novo peptide match those in the ground-truth. The reason is that a de novo peptide may often contain errors such as swapping or substitution of isobaric amino acid combinations, thus it may not exactly match the ground-truth but most of its predicted amino acids are still correct. The numbers of matched peptides and amino acids are summed up over all input spectra and then averaged by the number of ground-truth peptides and amino acids to obtain the peptide and amino acid accuracies, respectively. In this study, we did not apply any filter on the de novo results and all tools were expected to recover as many ground-truth peptides as they could.

We proposed a new metric named fragment ion accuracy and its motivation is as follows. Deep learning models are capable of learning many features, including some beyond the information provided by the mass spectra, such as the frequency or the sequential patterns of amino acids in the training data. While those extra features are helpful, and sometimes very powerful, we argue that a de novo peptide predicted from a tandem mass spectrum must have sufficient supporting evidence from that spectrum, i.e. the supporting fragment ions. Thus, when comparing a de novo peptide to a ground-truth on a tandem mass spectrum, we also counted how many peaks in the spectrum matched the theoretical ions of both de novo and ground-truth peptides. The fragment ion accuracy was then calculated as the percentage of theoretical ions of the ground-truth that were observed in both the de novo peptide and the spectrum. We believe this new metric provides a fair evaluation of de novo peptide sequencing tools to identify relevant fragment ions from the tandem mass spectra.

#### FDR estimation for de novo peptide sequencing

The three accuracy metrics above, however, still rely on a ground-truth peptide for every tandem mass spectrum. As mentioned in the introduction, the ground-truth peptides and PSMs are often obtained from the database search results, which may be biased towards high-quality spectra and well-known protein databases. More importantly, those metrics cannot measure how significant a de novo peptide-spectrum match is compared to a random match. Thus, an FDR estimation is desired to evaluate the de novo results.

In the database search approach, decoy peptides can be generated by randomly shuffling or reversing the target protein database. However, since most de novo peptide sequencing methods are designed to find the optimum amino acid sequence for a given tandem mass spectrum, a random decoy peptide is not likely to yield a better match to that spectrum. Thus, we decided to use decoy spectra rather than decoy peptides to establish an FDR evaluation for de novo peptide sequencing.

Given an input dataset of tandem mass spectra, here named “target spectra”, we generated a “decoy” spectrum for every “target” spectrum as follows. We started with a target spectrum, removed a number of its peaks, named “target peaks”, and then added back the same number of “noise peaks” to create a decoy spectrum. Firstly, the decoy spectrum has the same number of peaks as its target spectrum. Secondly, to create noise peaks, a simple way is to randomly generate their m/z and intensities from uniform distributions. However, the distributions of decoy spectra then may be very different from those of target spectra. Thus, we calculated the empirical m/z and intensity distributions of all the peaks in the target spectra and then randomly generated noise peaks from those empirical distributions. As a result, the m/z and intensity distributions of target peaks and noise peaks in the decoy spectra were very similar (Fig. 1a).

**Fig. 1.**
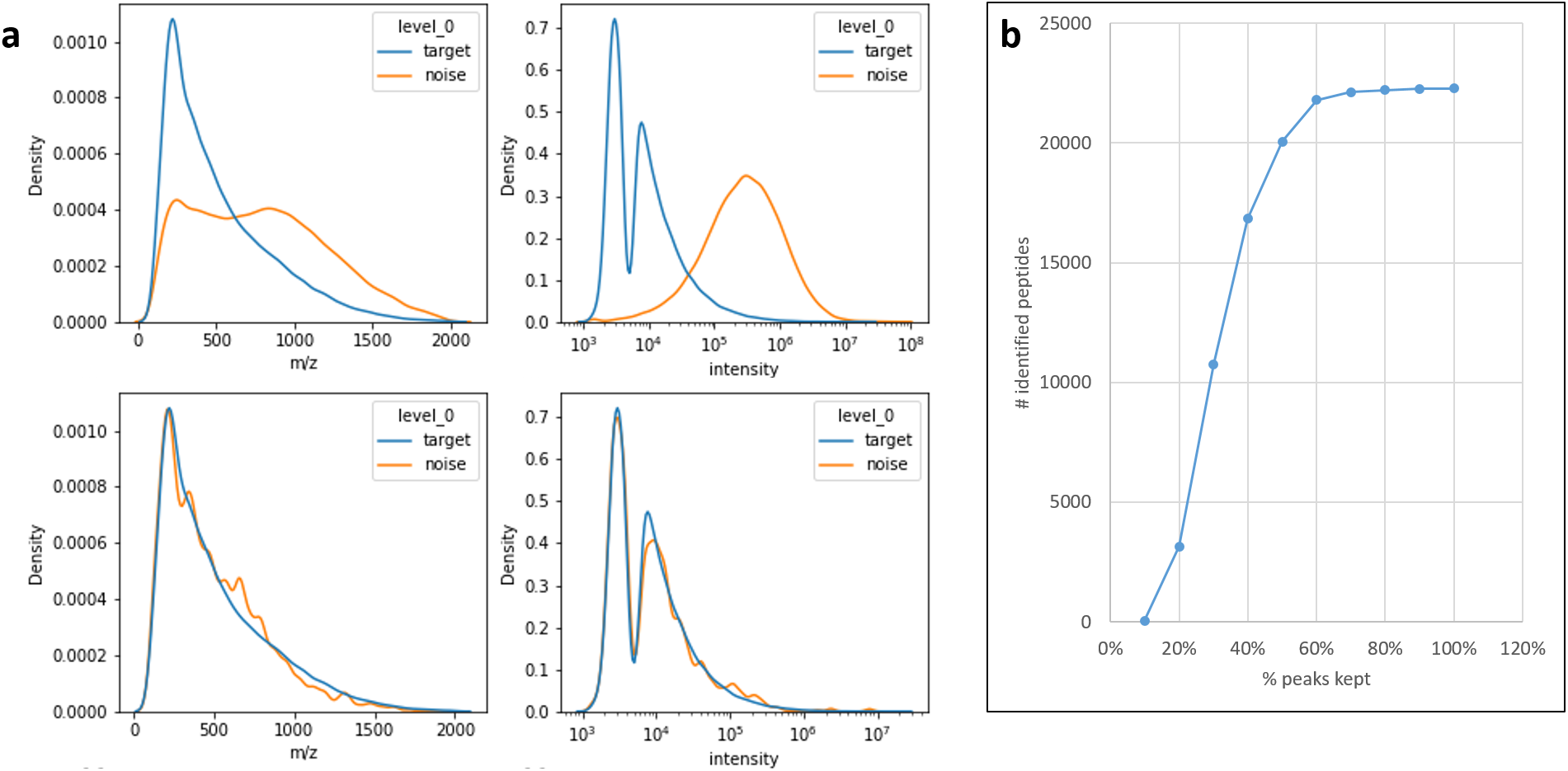
**a**, m/z and intensity distributions of the target and noise peaks in the decoy spectra generated from the ABRF dataset, where the noise peaks were randomly drawn from the uniform distributions (upper panels) and from the ABRF target spectra’s distributions (lower panels). **b**, Number of identified peptides from the database search of the decoy spectra decreased as the number of target peaks was reduced.

Finally, to determine the x% of target peaks to keep in a decoy spectrum, we experimented with different x from 0-100% to generate decoy spectra for the ABRF dataset and then performed a database search on those decoy spectra. We found that when x dropped below 10%, the database search could no longer identify any peptide (Fig. 1b). Thus, in this study, we kept 10% of the target peaks in a decoy spectrum. For example, given an input target spectrum with 100 peaks, we first randomly selected 10 peaks to keep and removed the other 90; then we added 90 noise peaks of which m/z and intensities were drawn from the empirical distributions of all the peaks in all the target spectra.

Since a target spectrum contains fragment ions of a real peptide whereas a decoy spectrum simply contains random peaks, a de novo sequencing tool is expected to predict a meaningful peptide sequence with a high-confidence score from a target spectrum, and an arbitrary sequence with a low score from a decoy spectrum. As a result, we could compare the de novo sequencing score distributions of the target spectra versus the decoy spectra to gain some insights, for example, how significant a target de novo PSM is compared to a decoy de novo PSM. Fig. 2 shows such an example of PEAKS, PointNovo, Casanovo, and GraphNovo on the ABRF dataset. A de novo FDR can also be calculated as the percentage of decoy de novo PSMs with scores above a given threshold. However, it is important to emphasize that a target de novo PSM here means that the de novo peptide matches the target spectrum with a score more significant than random matches, and it does not imply that the de novo peptide is the real peptide, as opposed to a target PSM in the database search approach. In other words, even when a de novo PSM has a significantly high score, the de novo peptide may still contain sequencing errors. Thus, the de novo FDR estimation here should be considered as a lower bound of the true FDR.

**Fig. 2.**
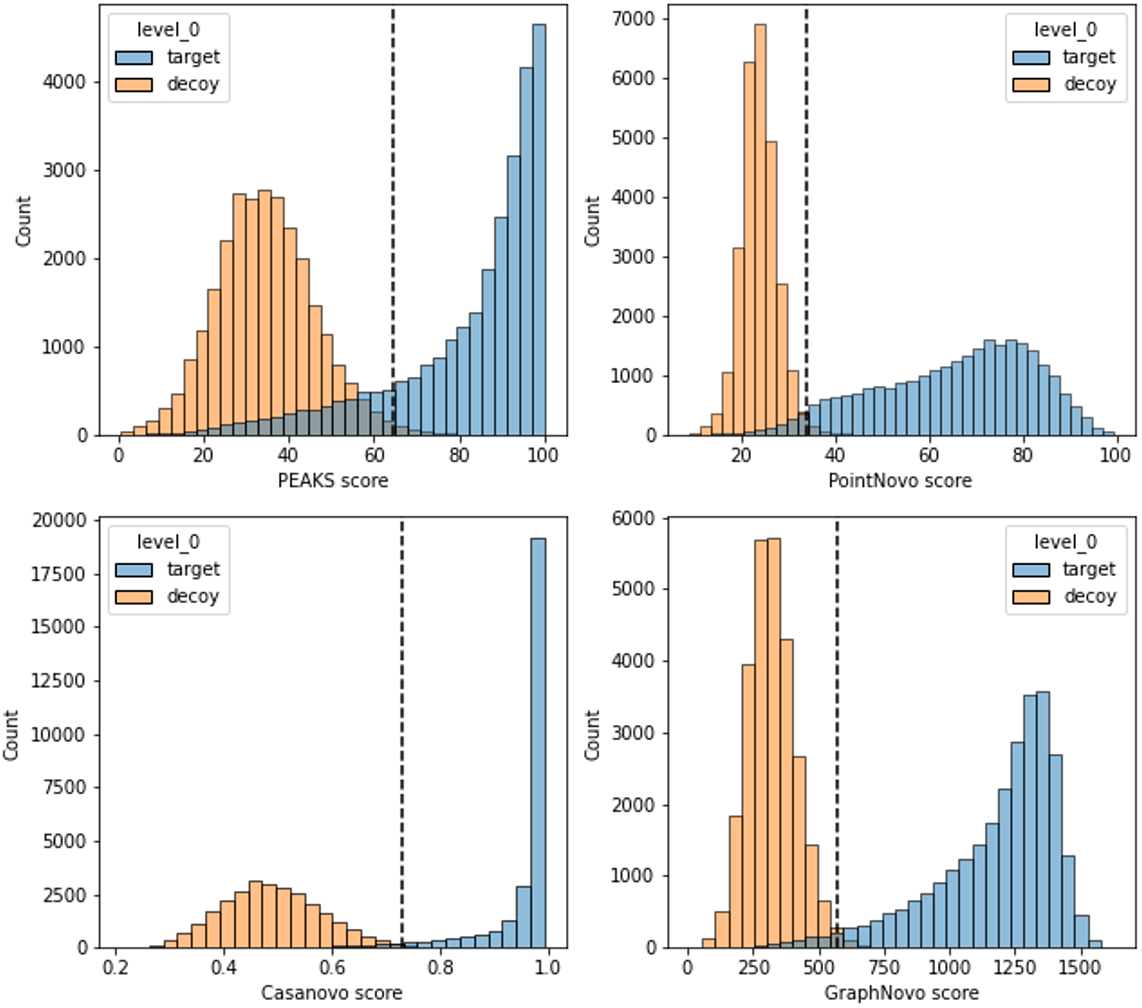
Distributions of the de novo sequencing scores on the tandem mass spectra of the ABRF dataset (blue) and their decoy spectra (orange). The dashed lines indicate the 1% FDR cutoff.

Another useful application of the FDR estimation and decoy spectra is to validate whether a de novo peptide sequencing tool tends to memorize peptide sequences. In particular, a decoy spectrum contains random peaks that are not likely to form a series of fragment ions of a peptide sequence. However, by memorizing the peptide sequences from the training, a de novo peptide sequencing tool may still can predict some sequence with a high score without supporting evidence from the decoy spectrum. Thus, we could evaluate the de novo peptide sequencing tools on their capability to differentiate between target and decoy spectra.

### Performance evaluation on the ABRF dataset

We first evaluated the performance of the four de novo peptide sequencing tools on the dataset from the 2017 ABRF PRG study (ABRF dataset). All three models PointNovo, Casanovo, and GraphNovo were trained on the MassIVE-KB human spectral library, which contains mostly tryptic peptides. Since the ABRF dataset is also human tryptic data, all four tools performed well on this dataset, with peptide, amino acid, and fragment ion accuracies at very high ranges of 43-76%, 74-88%, and 89-96%, respectively (Fig. 3a). The three deep learning tools performed better than PEAKS as expected, especially on the peptide accuracy. On the amino acid and fragment ion metrics, GraphNovo outperformed the other tools, thanks to its intensive efforts in modeling the features of fragment ions, i.e. the nodes in a spectrum graph^14^. Casanovo achieved a surprisingly high peptide accuracy of 76%, while its amino acid and fragment ion accuracies were 4-8% lower than GraphNovo. This discrepancy suggests that Casanovo might perform exceptionally well on some peptides and predict the whole sequences correctly, but for some other peptides, it might not be able to predict any correct amino acid.

**Fig. 3.**
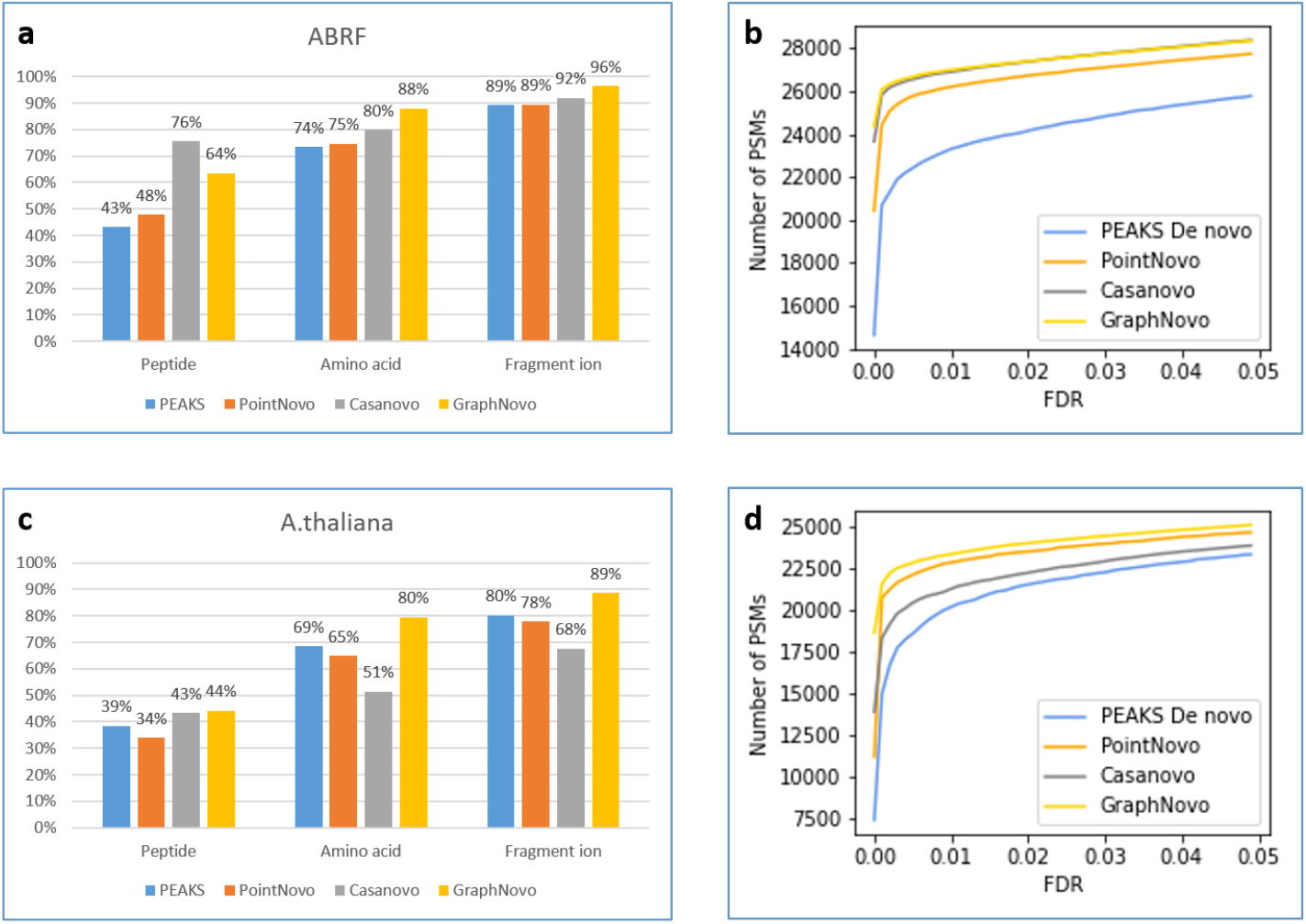
Evaluation of the de novo sequencing results on the ABRF **(a, b)** and the A. thaliana dataset **(c, d)**.

We further performed the FDR evaluation of the four de novo sequencing tools on the decoy spectra generated from the ABRF dataset. The score distributions of target and decoy de novo PSMs are shown in Fig. 2. We also calculated the de novo FDR as described above and reported the number of target de novo PSMs at different FDR cutoffs from 1-5% in Fig. 3b. For instance, at 1% FDR cutoff, PEAKS, PointNovo, Casanovo, and PointNovo reported 23,300, 26,194, 26,892, and 26,982 target de novo PSMs, respectively. The results in Fig. 2 and Fig. 3b show that GraphNovo and Casanovo were able to better differentiate between target and decoy spectra and hence reported more target de novo PSMs than PointNovo and PEAKS.

### Performance evaluation on the A. thaliana dataset

We next evaluated the performance of the four de novo peptide sequencing tools on a tryptic dataset from *A. thaliana* (PXD013658, Muntel et al.^7^). *A. thaliana* is a species that is less related to human and to the best of our knowledge, it has not been used to train any de novo sequencing models. As expected, the accuracies of the de novo results dropped substantially when compared to the ABRF dataset, with peptide, amino acid, and fragment ion accuracies at 34-44%, 51-80%, and 68-89%, respectively (Fig. 3c, d). While the performance drop was expected as the models were trained on human data, not *A. thaliana*, Casanovo suffered considerably. Its peptide accuracy decreased by 33% (76% to 43%) while its amino acid and fragment ion accuracies even fell behind PEAKS and PointNovo. GraphNovo outperformed the other three tools on all three accuracy metrics and also on the FDR evaluation. The results on this *A. thaliana* dataset represent a good validation test showing that deep learning models are biased towards the training data to different extents. It is also interesting to see that PEAKS performance was quite robust, as its peptide and amino acid accuracies only dropped around 4-5%, much less than the other three deep learning tools.

### Performance evaluation on the nontryptic and the HLA datasets

Since tryptic peptides are over-represented in most public MS data repositories, including MassIVE-KB, de novo peptide sequencing tools may be biased towards those peptides. Thus, it is essential to assess how they would perform on nontryptic data. Here we performed evaluation on a dataset from Wang et al.^6^ (PXD010154) that includes peptides digested from six enzymes: Arg-C, Asp-N, Chymotrypsin, Glu-C, Lys-C, and Lys-N. Casanovo has a nontryptic model that was fine-tuned on their MassIVE-KB model using the PSMs of peptides with diverse C-terminal amino acids obtained from MassIVE-KB and PROSPECT databases^9,12,19^. PointNovo has two models, one trained on MassIVE-KB and another trained on HLA data; we tested both and reported the results of the HLA model here since it showed better performance. For GraphNovo and PEAKS, we still used the same models as before because they do not have other options.

The nontryptic evaluation results are shown in Fig. 4a, b. The overall performance of all tools are considerably lower than the previous results on tryptic data (Fig. 3). GraphNovo outperformed the other tools by a large margin; its peptide, amino acid, and fragment ion accuracies were 6%, 14%, and 10%, respectively, higher than the second-best tools. This is despite the fact that GraphNovo model was trained on tryptic data, whereas PointNovo and Casanovo models had been fine-tuned on nontryptic data. The robust performance of GraphNovo over the other tools was also confirmed in the FDR evaluation (Fig. 4b).

**Fig. 4.**
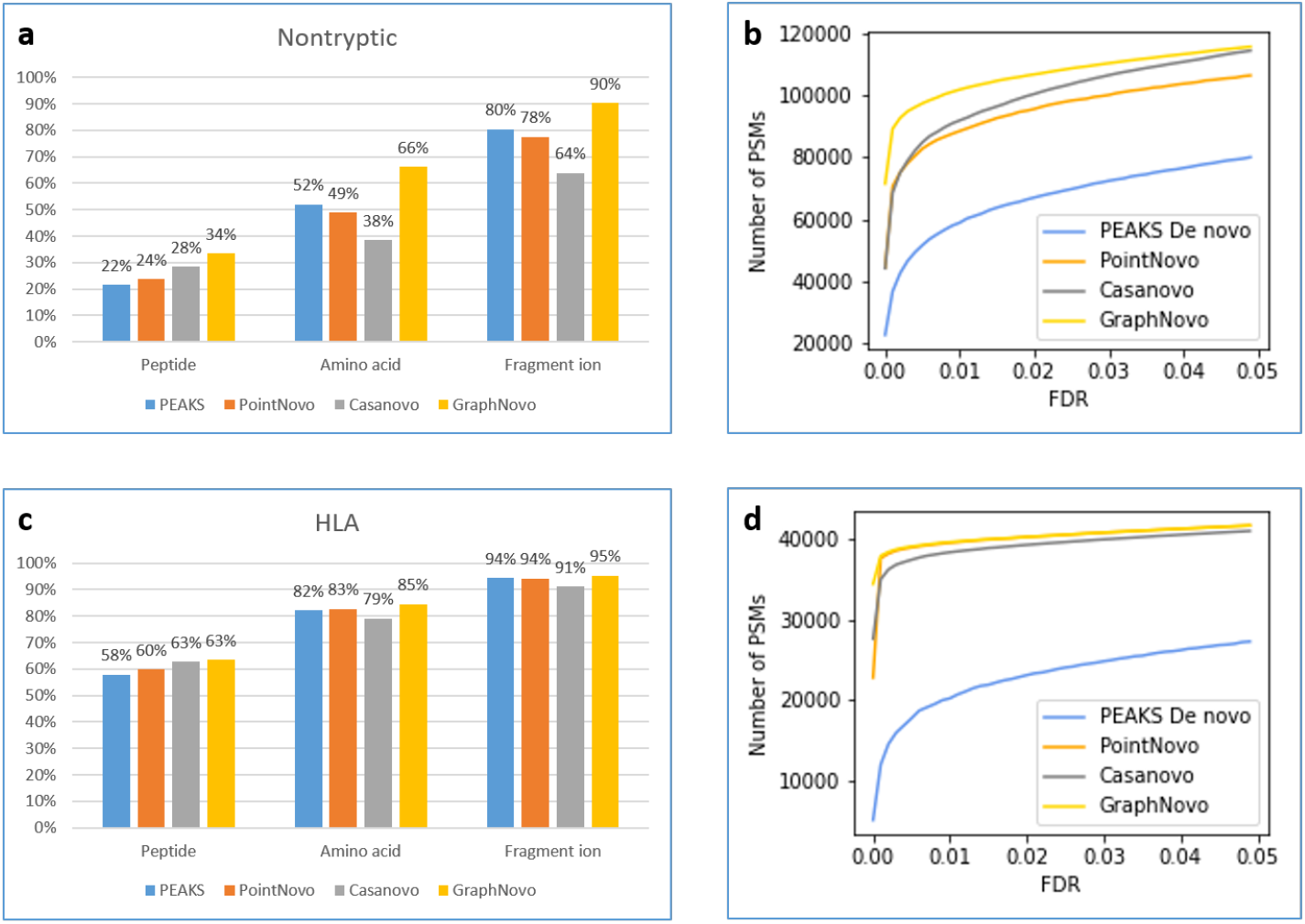
Evaluation of the de novo sequencing results on the nontryptic **(a, b)** and the HLA dataset **(c, d)**.

We further performed evaluation on an HLA class 1 dataset (PXD021013) from Wilhelm et al.^8^. HLA peptides are also very different from tryptic peptides due to their unspecific digestion and other unique characteristics such as length and binding motifs. As shown in Fig. 4c, all four tools performed well on this dataset with very high peptide accuracy from 58-63%. This is probably because HLA class 1 peptides are short, with lengths from 8-12 amino acids, hence it is easier for the de novo peptide sequencing tools to predict the whole peptide sequences correctly.

There was no significant difference between the accuracies of the four tools. On the FDR evaluation, GraphNovo and PointNovo reported the most number of target de novo PSMs, followed closely by Casanovo.

### Performance evaluation unseen mutated peptides

Finally, we evaluated the de novo sequencing methods on the dataset of ten thousand unseen mutated peptides, which were originally selected from the MassIVE-KB database^9^ and then randomly mutated by 1-10 amino acids. Their spectra were predicted using the 2020 HCD model of Prosit^8^. For PointNovo, Casanovo, and GraphNovo, we used their models that were trained on the MassIVE-KB database. By starting with MassIVE-KB peptides and gradually increasing the number of mutations, we can investigate the impact of mutations on the performance of each tool.

Fig. 5 shows the peptide, amino acid, and fragment ion accuracies of the de novo results with respect to the number of mutations. As expected, the overall performance decreased as the number of mutations increased from 0 to 10. On the peptide metric, Casanovo showed the steepest decrease of 21% (from 45% to 24%), followed by GraphNovo (from 33% to 23%) and PEAKS (from 16% to 10%), while PointNovo only showed minor decline (from 22% to 19%). On the amino acid metric, GraphNovo and PointNovo showed a modest decrease of about 8%, while Casanovo and PEAKS dropped by about 20%. On the fragment ion metric, GraphNovo and PointNovo also showed very little change, while Casanovo dropped by about 16%. Overall, Casanovo seems to be the most sensitive to the number of mutations, followed by PEAKS, GraphNovo, and PointNovo. We also observed a similar trend to the previous datasets that Casanovo had higher peptide accuracy, but fell behind the other tools on the amino acid and fragment ion accuracies.

**Fig. 5.**
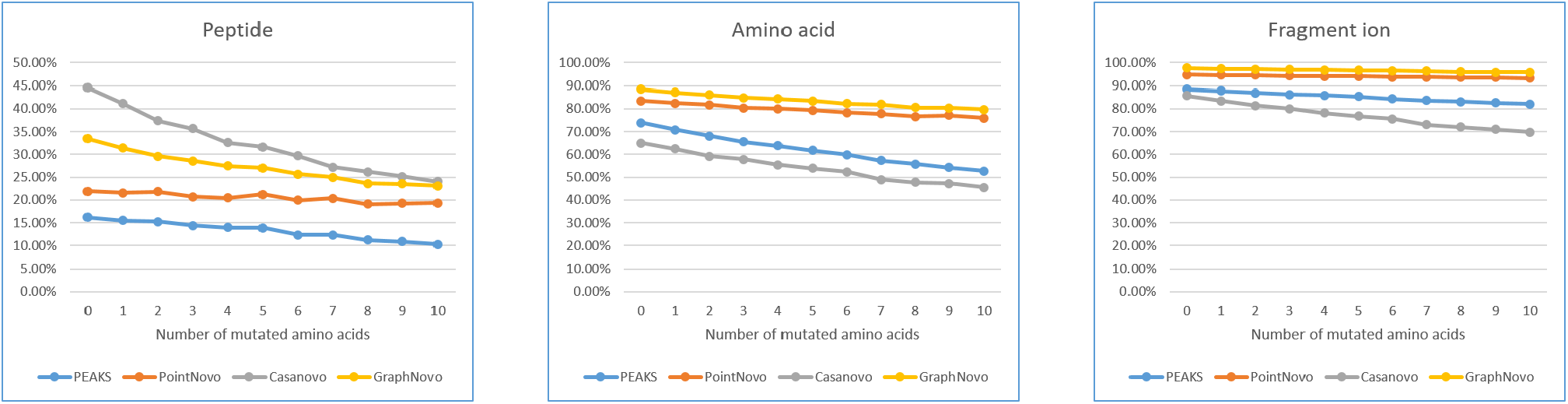
Evaluation of the de novo sequencing results on the simulated dataset of unseen mutated peptides. **a**, Peptide accuracy. **b**, Amino acid accuracy. **c**, Fragmention accuracy.

## Discussion

In this study, we proposed NovoBoard, a comprehensive framework to evaluate the performance of de novo peptide sequencing methods. The framework aims to establish a standard set of benchmark datasets and metrics that can thoroughly assess the strengths and weaknesses of de novo peptide sequencing methods and also their specific applications.

Special focuses were put into addressing deep learning-based models to validate that they were truly able to generalize and discover new sequences, rather than overfitting and memorizing the training data. We also tried to propose an FDR estimation based on decoy spectra for de novo peptide sequencing, which can be used to assess how significant a de novo peptide-spectrum match is compared to a random match. Overall, the results in our study indeed revealed important characteristics of different de novo peptide sequencing methods that will be valuable for future applications. The results also confirmed that such a comprehensive evaluation framework is crucial and needs more attention from the research community.

One should note again that it is always possible that a de novo peptide may contain sequencing errors regardless of how good its PSM and supporting fragment ions are. This is in contrast to a database peptide, which comes from a known protein database. Thus, our de novo FDR estimation here should be considered as a lower bound of the true FDR.

Another limitation of our study is that the benchmark datasets mainly came from Orbitrap instruments and DDA experiments. Thus, they could be extended to include data from other MS instruments, and especially, from DIA experiments. However, it should also be noted that de novo peptide sequencing methods for DIA data are not very popular and their evaluation may be more complicated than for DDA data.

In this study we did not perform a detailed analysis on the computing time and resources of the de novo sequencing tools. We believe that it is more related to the engineering efforts and the available hardwares. Nevertheless, we did a quick test on the ABRF dataset and reported here a rough estimation of the tools’ running time: PEAKS 100 spectra/second (on CPU i9-13900K), PointNovo 110 spectra/second, Casanovo 150 spectra/second, and GraphNovo 273 spectra/second (the later three were run on GPU, NVIDIA GeForce RTX 4090). It is worth noting that a speed of >=250 spectra/second should be able to keep up with the data acquisition speed of most MS instruments and enable real-time de novo sequencing directly on the instruments.

## Methods

### Datasets and search parameters

The ABRF dataset was obtained from the 2017 Association of Biomolecular Resource Facilities (ABRF) Proteomics Research Group (PRG) study. The nontryptic dataset was obtained from Wang et al.^6^ (PXD010154). The *A. thaliana* dataset was obtained from Muntel et al.^7^ (PXD013658). The HLA class 1 dataset was obtained from Wilhelm et al.^8^ (PXD021013). We also simulated a dataset of randomly mutated peptides and their *in silico* spectra. First, ten thousand peptides were randomly selected from the MassIVE-KB database^9^, then 1-10 amino acid mutations were introduced to each peptide and finally their spectra were predicted using Prosit^8^.

The raw data of all five datasets were converted to MGF files using the Proteome Discoverer software. We also performed database search of those datasets using Proteome Discoverer with the following parameters: precursor mass tolerance 10 ppm, fragment ion mass tolerance 0.02 Da, fixed Cysteine(Carbamidomethylation), variable Methionine(Oxidation), human and *Arabidopsis thaliana* protein databases (version SwissProt 20240224), and peptide FDR 1%. PEAKS, PointNovo, Casanovo, and GraphNovo were run on the MGF files. The de novo sequencing parameters include precursor mass tolerance 10 ppm, fragment ion mass tolerance 0.02 Da, fixed Cysteine(Carbamidomethylation), and variable Methionine(Oxidation). No filter was set on the output de novo results.

PEAKS and GraphNovo each have only one single model which was applied to all five datasets. PointNovo has a tryptic model and an HLA model; the tryptic model was applied to the ABRF, the A. thaliana, and the mutated datasets; the HLA model was applied to the nontryptic and the HLA datasets. Casanovo has a tryptic model and a nontryptic model; the tryptic model was applied to the ABRF, the A. thaliana, and the mutated datasets; the nontryptic model was applied to the nontryptic and the HLA datasets.

PEAKS, PointNovo, Casanovo, and GraphNovo were run on a machine equipped with an Intel CPU (i9-13900K), an NVIDIA GPU (GeForce RTX 4090), and 128 GB memory. PEAKS, PointNovo, and GraphNovo were implemented and run in PEAKS Studio in Windows environment, while Casanovo was run in Linux environment.

## Data availability

The raw data of the datasets can be obtained from the ProteomeXchange Consortium via the PRIDE^20^ partner repository with the dataset identifiers PXD010154, PXD013658, and PXD021013.

## Acknowledgements

We thank Dr. William Stafford Noble for his advice on setting the appropriate parameters for Casanovo

## Author contributions

Ngoc Hieu Tran performed the evaluation analysis. Rui Qiao proposed the idea of FDR estimation for de novo peptide sequencing. Zeping Mao designed the GraphNovo model. Shengying Pan implemented the GraphNovo software. Qing Zhang and Wenting Li prepared the benchmark datasets. Lei Xin and Baozhen Shan proposed and planned the study. Lei Xin, Baozhen Shan, and Ming Li supervised the study.

